# Functional connectomics of affective and psychotic pathology

**DOI:** 10.1101/489377

**Authors:** Justin T. Baker, Daniel G. Dillon, Lauren M. Patrick, Joshua L. Roffman, Roscoe O. Brady, Diego A. Pizzagalli, Dost Öngür, Avram J. Holmes

**Affiliations:** McLean Hospital, Belmont, MA 02478; Department of Psychiatry, Harvard Medical School, Boston, MA 02114; Department of Psychology, Yale University, New Haven, CT 06520; Athinoula A. Martinos Center for Biomedical Imaging, Charlestown, MA 02129; Massachusetts General Hospital, Boston, MA 02114; Department of Psychiatry, Beth Israel Deaconess Medical Center, Boston, MA 02114; Department of Psychiatry, Yale University, New Haven, CT 06520

## Abstract

Converging evidence indicates that groups of patients with nominally distinct psychiatric diagnoses are not separated by sharp or discontinuous neurobiological boundaries. In healthy populations, individual differences in behavior are reflected in variability across the collective set of functional brain connections (functional connectome). These data suggest that the spectra of transdiagnostic symptom profiles observed in psychiatric patients may map onto detectable patterns of network function. To examine the manner through which neurobiological variation might underlie clinical presentation we obtained functional magnetic resonance imaging (fMRI) data from over 1,000 individuals, including 210 diagnosed with a primary psychotic disorder or affective psychosis (bipolar disorder with psychosis and schizophrenia or schizoaffective disorder), 192 presenting with a primary affective disorder without psychosis (unipolar depression, bipolar disorder without psychosis), and 608 demographically and data-quality matched healthy comparison participants recruited through a large-scale study of brain imaging and genetics. Here, we examine variation in functional connectomes across psychiatric diagnoses, finding striking evidence for disease connectomic “fingerprints” that are commonly disrupted across distinct forms of pathology and appear to scale as a function of illness severity. Conversely, other properties of network connectivity were preferentially disrupted in patients with psychotic illness, but not patients without psychotic symptoms. This work allows us to establish key biological and clinical features of the functional connectomes of severe mental disease.

**SIGNIFICANCE STATEMENT:** Historically, most research on the biological origins of psychiatric illness has focused on individual diagnostic categories, studied in isolation. Mounting evidence suggests nominally distinct psychiatric diagnoses are not separated by clear neurobiological boundaries. Here, we derive functional connectomic signatures in over 1,000 individuals including patients presenting with specific categories of impairment (psychosis), clinical diagnoses, or severity of illness as reflected in treatment seeking. Our analyses revealed features of connectome functioning that are commonly disrupted across distinct forms of pathology, scaling with clinical severity. Conversely, other aspects of network connectivity were preferentially disrupted in patients with psychotic illness, but not patients without psychotic symptoms. These data have important implications for the establishment of functional connectome fingerprints of severe mental disease.

## INTRODUCTION

Recent progress in the neurosciences has provided unprecedented opportunities for advancing our understanding of the etiology and pathogenesis of psychiatric illness. At the same time, the gradual reification of diagnostic categories has hampered our ability to take full advantage of these innovations^1–4^. To date, the vast majority of research on the biological origins of psychopathology has focused on discrete illness categories, studied in isolation. Although modern psychiatric diagnoses provide advantages to the field in terms of diagnostic reliability, their construct validity and utility for understanding brain circuit dysfunction has been challenged^2,3^. Converging epidemiologic, genetic, and neuroscientific research suggests that populations of psychiatric patients are not separated by clear neurobiological borders between diagnostic categories or across health and disease. There is evidence, for instance, of substantial overlap in the genetic factors that increase risk for both affective and psychotic illness^5–7^. Consistent with shared heritability, partially overlapping patterns of brain network dysfunction mark a broad range of mental diseases^8–10^, indicating that their breakdown can lead to diverse forms of psychopathology. Yet, despite a flurry of important scientific advances, we still remain far from a mechanistic understanding of how the functioning of large-scale brain networks might serve to influence suites of behaviors, within, or across psychiatric illnesses.

Identifying signatures of pathology across the functional connectome could provide a framework for researchers to study neurobiological contributions to the onset and maintenance of clinically relevant symptoms, informing the development of novel treatments and future classification schemes. Emerging evidence in healthy populations suggests that individual differences in behavior may be reflected in variability across the collective set of functional brain connections^11–13^ (functional connectome)^14^. Work from our group and others indicates that the unique connectome architecture of an individual’s brain serves as a stable and reliable “fingerprint”^12,13,15–17^, likely influenced by genetic variation^18–20^. The spectra of symptom profiles observed in patient populations may arise through detectable patterns of network function^1,21,22^. In particular, the disturbance of individual networks might preferentially contribute to domain-specific (e.g., executive, affective, and social), but disorder-general, impairments^8,9,21^. While some common patterns of network functioning may be shared across illnesses, for example the hypothesized central role of altered frontoparietal network connectivity in mental health^8^, other network-specific alterations may produce clusters of symptoms that preferentially present in specific illnesses (e.g., psychosis).

Despite increasing interest in the study of relations that link connectome functioning with broad diagnostic syndromes, existing work in this domain is often limited in several key respects. First, most research utilizes case-control designs, examining single psychiatric illnesses in isolation. This approach can potentially mask the presence of substantial overlap in the distributions of connectome functioning across populations, providing the illusion of group specificity. Complex clinical phenotypes arise from coordinated interactions throughout the functional connectome^11^. High co-morbidity across illnesses suggests the presence of dimensional network-level abnormalities that bridge across traditional diagnostic constructs^10^. To achieve a breakthrough in our understanding of how brain functions underlie psychiatric illness, we must collect datasets that span diagnostic categories. Second, prior research on the connectomic signatures of psychiatric illness has largely focused on circumscribed patient samples recruited either from single clinical settings *<u>or</u>* the broader community. As a consequence, we are often unable to assess the manner in which the connectome associates of psychiatric illness may vary as a function of symptom severity, for instance, as indicated by degree of treatment seeking.

Leveraging this connectome approach, we recently identified disturbances within the frontoparietal control network (spanning aspects of dorsolateral prefrontal, dorsomedial prefrontal, lateral parietal, and posterior temporal cortices) in patients with schizophrenia and psychotic bipolar disorder^23^. Impairments in the integration and processing of information across large-scale brain networks are thought to mark psychotic illness^24^. Our prior work, for instance, revealed frontoparietal network abnormalities, preferentially evident in the control B subnetwork, in patients diagnosed with schizophrenia and bipolar disorder with psychotic features^23^. Higher-order task-activations recruit cortical territories in the frontoparietal network^25^, with the highest-order task responses being most consistent within the control B subnetwork. While frontoparietal network disruption could underlie a specific vulnerability for thought disorder that characterizes psychosis, there is evidence for functional alterations in this system across a range of patient populations^8^, including unipolar depression, bipolar disorder and schizophrenia^23,26^. For example, regional impairments within aspects of the frontoparietal network are thought to contribute to both depressive^27^ and manic episodes^28^, as well as the occurrence of psychotic symptoms^29^. Despite these converging lines of evidence, the extent to which dysfunction in frontoparietal connectivity tracks the presence of specific diagnostic categories (e.g., psychotic illnesses), symptom severity, utilization of care, or other unmeasured factors is poorly understood.

In the current study we investigate whether patterns of connectomic disruption in psychiatric illness track specific categories of impairment (presence of psychosis), clinical diagnoses (unipolar depression, bipolar disorder, and schizophrenia or schizoaffective disorder), or severity of illness as reflected in treatment seeking. First, we demonstrate that both affective illness without psychosis and psychotic illness broadly associate with reduced connectivity across multiple large-scale cortical networks. Consistent with the hypothesized central role of disrupted executive functions in mental health, our analyses reveal a graded pattern of dysconnectivity within the frontoparietal network, which is amplified in patients suffering from more extreme forms of psychopathology. This transdiagnostic profile of impairment is present in patients with and without psychotic symptoms and across individual diagnostic categories. Suggesting a link between disrupted frontoparietal network functioning and diagnostic severity, loss of connectivity is evident in treatment seeking patients with unipolar depression, but not within non-treatment seeking individuals who met criteria for unipolar depression recruited from the general community. Second, the observed patterns of connectome functioning display evidence of general as well as specific alterations in network connectivity across categories of impairment (psychosis) and clinical diagnoses. Schizophrenia and psychotic bipolar disorder, for instance, associate with a preferential reduction in default network integrity that is absent in affective illnesses without psychosis. Together, these results suggest that graded impairments within key control networks likely represent a common biological substrate central to the pathophysiology of both affective and psychotic illness, while other aspects of network function may preferentially link to specific symptom domains or diagnoses.

## RESULTS

Between November 2008 and June 2017, resting-state functional magnetic resonance imaging (fMRI) data were collected from 1,010 individuals, including 210 diagnosed with a primary psychotic disorder (137 meeting criteria for schizophrenia or schizoaffective disorder, 73 with bipolar disorder with psychosis), 192 presenting with a primary affective disorder without psychosis (26 with bipolar disorder without psychosis, 57 treatment seeking individuals with unipolar depression, 109 non-treatment seeking individuals with unipolar depression), and 608 demographically and data-quality matched healthy comparison participants recruited through an ongoing, large-scale study of brain imaging and genetics^30^ (Table S1). A subset of the participants recruited from McLean Hospital were included prior published analyses (60 meeting criteria for schizophrenia or schizoaffective disorder, 40 with bipolar disorder with psychosis)^23^.

To examine the functional network interactions affected by psychiatric illnesses with or without psychosis, we first processed the data with a series of steps common to intrinsic connectivity analyses^31–33^ and then computed cortical functional coupling matrices for each participant across all available parcels within the 17-network functional atlas of Yeo et al. 2011^34^. Additional details on the preprocessing procedures are detailed in Holmes et al., 2015^30^ and Yeo et al., 2011^34^. Next, we compared z-transformed Pearson correlation values across 3 groups (affective illnesses without psychosis, psychotic illness, healthy comparison participants) for all 3,660 (61×60) pairwise regional interactions (excluding unity interactions; Figure 1). Residual differences between the patient groups, relative to healthy comparison participants, are displayed in Figure 1E-F. Broad reductions in correlation between regions that spanned several functional networks, including the frontoparietal, default, and ventral attention networks were evident in patients with psychiatric illness, particularly those with psychosis. Across all intra-hemispheric cortical connections, 412 (11.26%) exhibited significant between group differences (FWER-corrected p≤0.05, corresponding to an uncorrected p≤1.37×10^-5^). When considering a less stringent statistical criterion (FDR-corrected q≤0.05, corresponding to an uncorrected p≤8.47×10^-4^) the number increased to 778 (21.26%). Between-network (i.e. off-diagonal) coupling was less negative, or muted, in the affective illnesses groups with and without psychosis (Figure 1E-F), consistent with a general flattening of intrinsic network connectivity across the connectome in patient populations.

**Figure 1.**
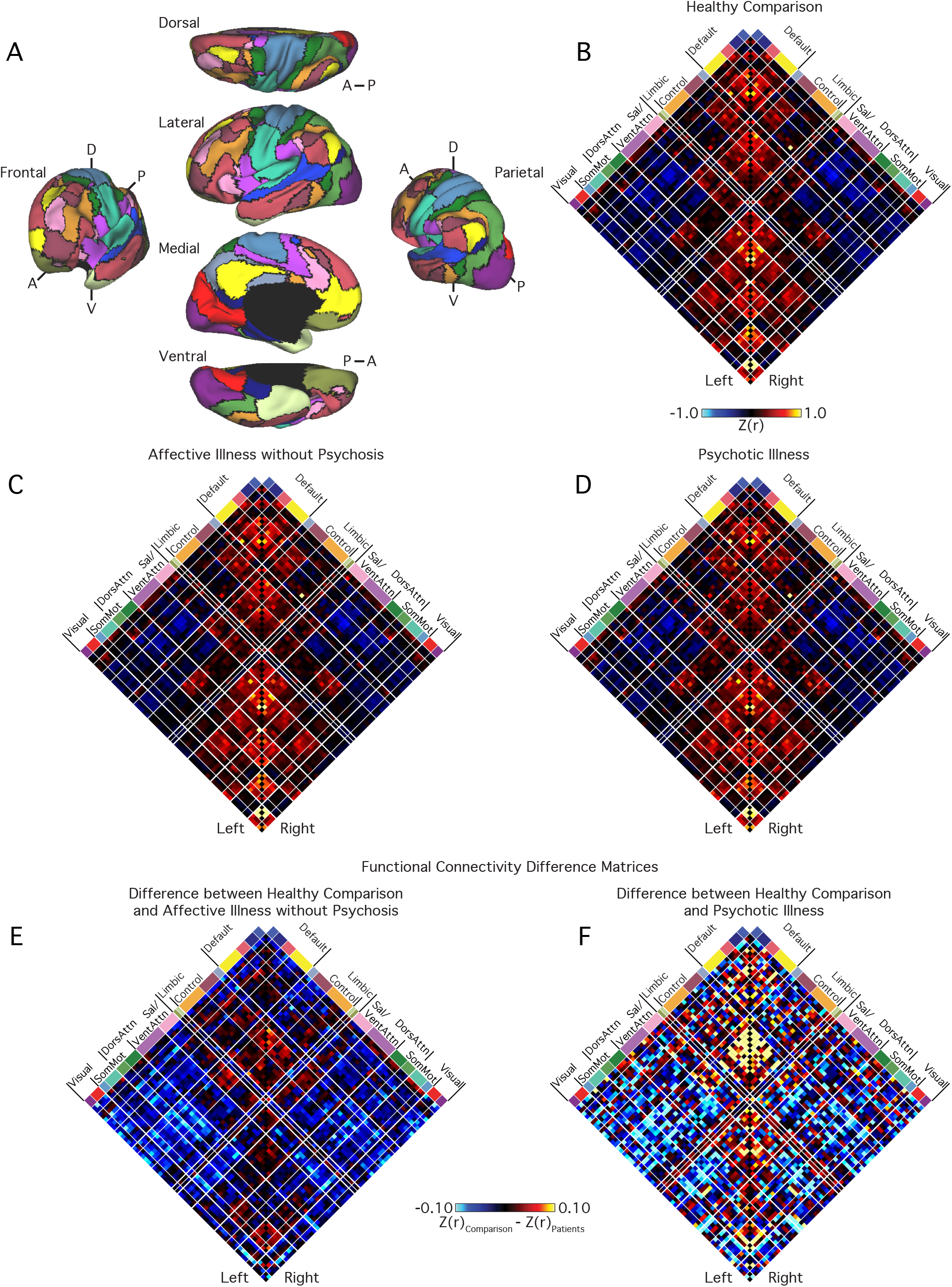
Cortical network connectivity in patients and healthy comparison participants. (A) The functional network organization of the human cerebral cortex revealed through intrinsic functional connectivity. Colors reflect regions estimated to be within the same network. Regions determined based on the 17-network solution from Yeo et al. 2011. The approach groups similar correlation profiles based on a winner-take-all solution, with every surface vertex assigned to its best-fitting network. The 2D grids (B-D) display the complete coupling architecture of the cerebral cortex measured at rest for (B) the healthy comparison participants, (C) patients with affective illnesses without psychosis, and (D) patients with psychotic illnesses. Values reflect z-transformed Pearson correlations between every region and every other region, after accounting for the effects of coil, scanner, console software version, age, sex, race, ethnicity, and handedness. Within-network correlations fall along the center diagonal. Between-network correlations are plotted away from the diagonal and reveal both positive (red) and negative (blue) correlations. (E-F) The 61×61 grids show the differences in resting blood oxygenation level–dependent (BOLD) correlation between controls and (E) patients with affective illnesses without psychosis, as well as (F) patients with psychotic illnesses, for each intra-hemispheric regional pair. Differences were obtained by an analysis of variance of z-transformed Pearson correlation values, adjusting for nuisance variables. White lines represent network boundaries. DorsAttn indicates dorsal attention; Left, left hemisphere; Right, right hemisphere; Sal, salience; SomMot, somatomotor; and VentAttn, ventral attention.

### Affective and psychotic illnesses associate with graded disruptions in frontoparietal network connectivity

In line with the core role of executive functioning deficits in mental health^35^, a growing literature suggests that frontoparietal network impairments may reflect a transdiagnostic marker of psychopathology^1,8,21^. We first tested the hypothesis that frontoparietal dysconnectivity would show a graded relationship with diagnostic status, increasing in severity in patients suffering from more extreme forms of psychopathology. In the frontoparietal network, we observed a main effect of *Group* (F_2,1001_=63.95, p≤0.001; μ^2^=0.11). Marked diagnosis-related differences in functional connectivity were evident for within-network connections involving the control B aspect of the frontoparietal network, with 90 percent of associated pairwise network combinations surviving corrections for multiple comparisons (18 of 20 region pairs; Figure 2A). Follow-up analyses revealed reduced mean control B network connectivity in both affective illnesses without psychosis (0.56±0.16; p≤0.01) and psychotic illness (0.42±0.18; p≤0.001) relative to the healthy comparison sample (0.58±0.16; Table S2). Furthermore, the connectivity of this aspect of the frontoparietal network was significantly reduced in patients with psychosis relative to those without psychotic symptoms, consistent with a graded reduction of network function (p≤0.001; Figure 3; see Table S2 in the Supplement for control A and B interactions).

**Figure 2.**
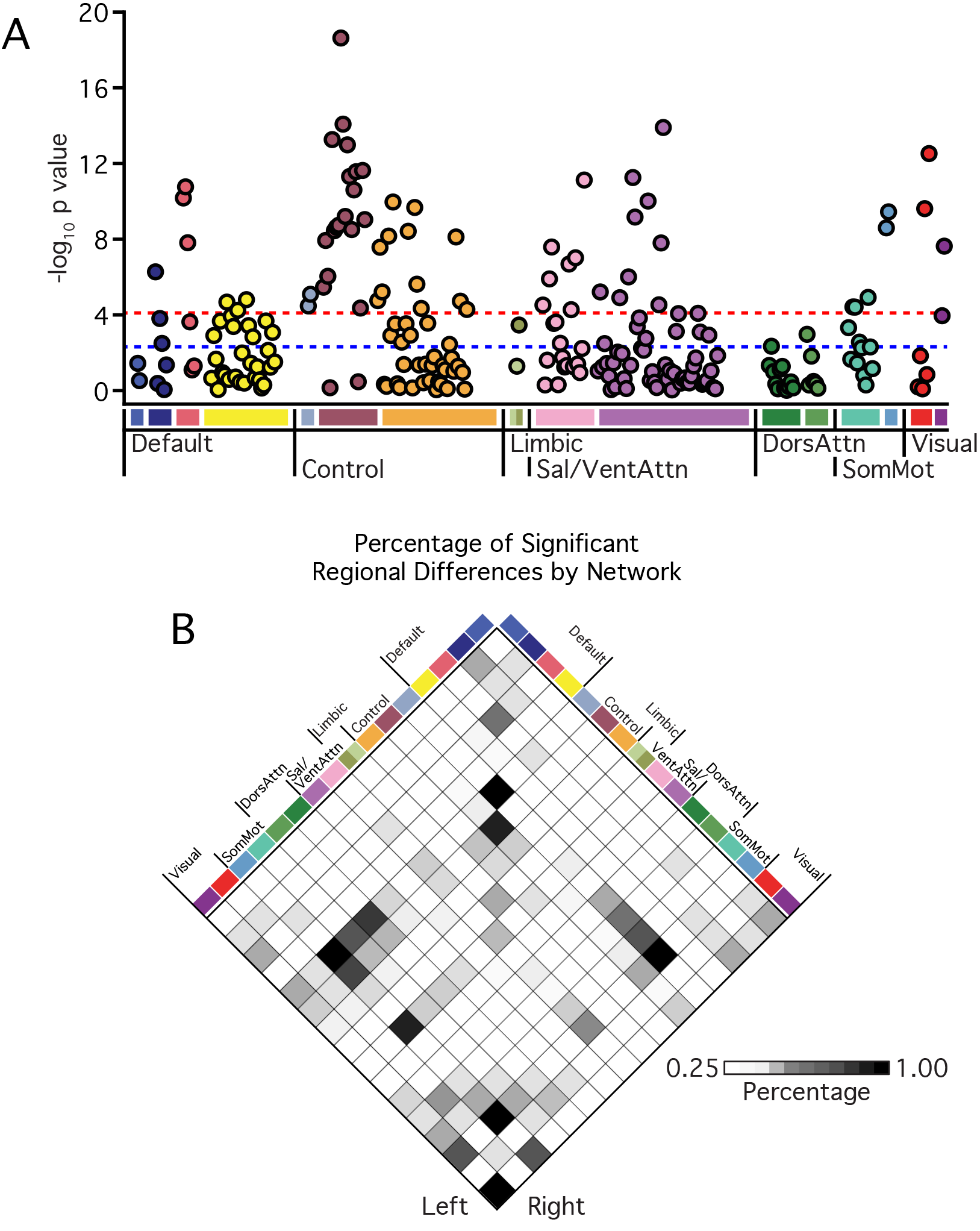
Affective illnesses without psychosis and psychotic illness broadly link to reductions in network connectivity across multiple functional networks. (A) Manhattan plot showing associated network-wide p-values of group-related (healthy comparison, affective illnesses without psychosis, psychotic illnesses) differences in functional connectivity. The y-axis shows the −log10 p-values of 240 within-network regional pairs, and the x-axis shows their network positions. The horizontal red line represents the threshold of p≤0.05 for Bonferroni-corrected significance across all possible regional pairs; the horizontal blue line represents the false discovery rate threshold of q≤0.05. (B) Each grid box represents the percentage of connections within and between networks that show a significant main effect of group at the threshold of p≤0.05 for Bonferroni-corrected significance across all possible regional pairs. See Figure 1 legend for explanation of abbreviations.

**Figure 3.**
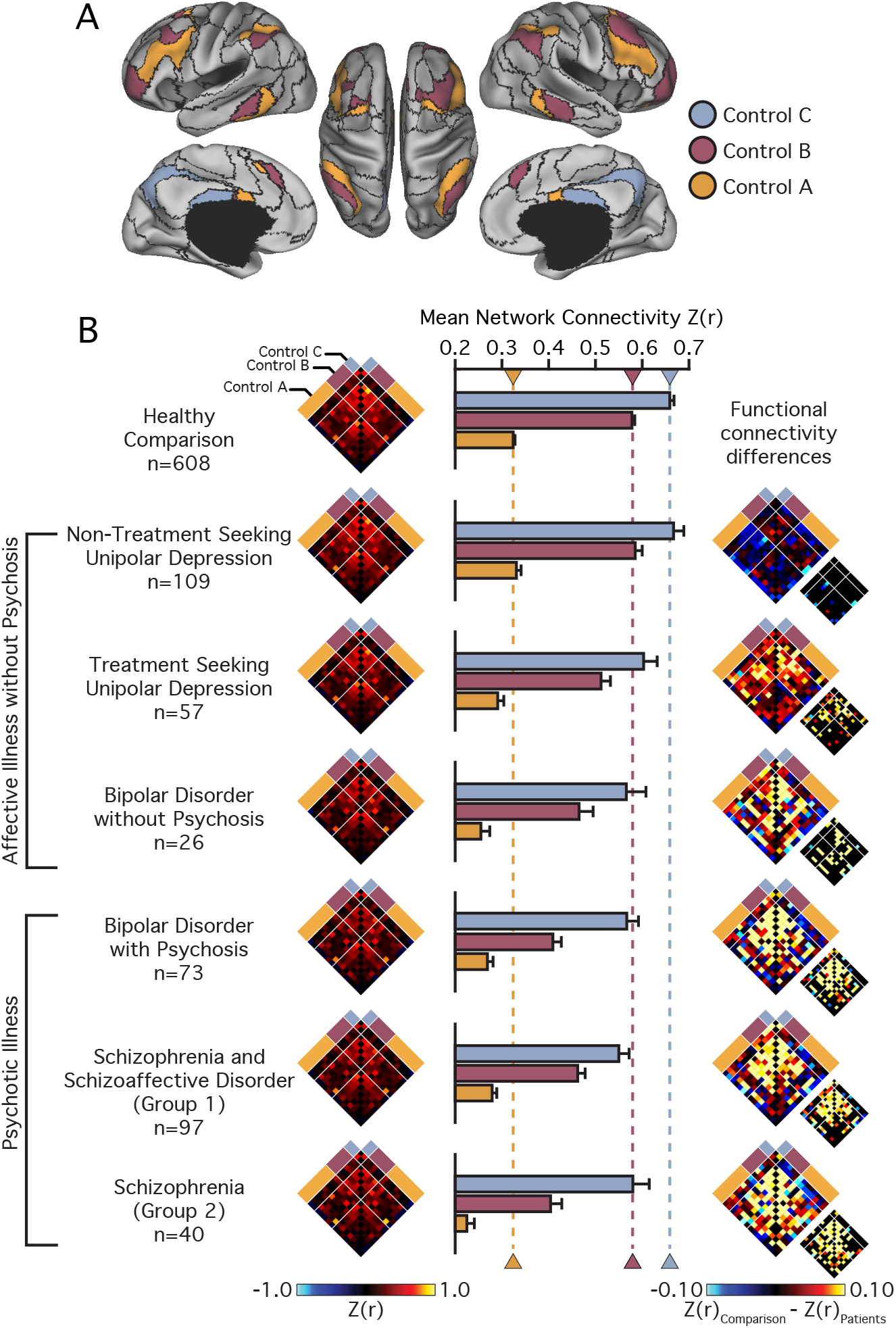
Disruptions in frontoparietal network connectivity increase with diagnostic severity. A) The colored aspects of cortex reflect regions estimated to be within the A, B, and C aspects of the frontoparietal control network. Black lines denote network boundaries defined through the 17-network solution from Yeo et al. 2011. (B) Functional connectivity matrices for the 14 left and right hemisphere regions of the frontoparietal control network shown for the healthy comparison and each patient group. Bar graphs reflect mean network connectivity for each group. Error bars denote standard error. Functional connectivity difference matrices were obtained by an analysis of variance of z-transformed Pearson correlation values, accounting for the effects of the coil, effects of scanner, console software version, age, sex, race, ethnicity, and handedness. Differences significant at false discovery rate (q≤0.05) are shown in each panel just to the lower right of the unthresholded matrix.

We next considered how this profile of frontoparietal network disruption may present across individual DSM-IV diagnoses. Participants with psychiatric illness were divided into non-treatment seeking individuals with unipolar depression (n=109), patients recruited from clinical services at McLean Hospital with unipolar depression (n=57), bipolar disorder without psychosis (n=26), and bipolar disorder with psychosis (n=73). The available sample of patients with schizophrenia was divided into two groups to examine the stability of observed effects across both samples and collection sites, this was not possible for other diagnostic groups, which were each recruited from a single clinical setting. Schizophrenia groups 1 (n=97) and 2 (n=40) were recruited through the respective inpatient services at McLean Hospital and Massachusetts General Hospital.

Analyses of region-to-region correlation strength across the frontoparietal network revealed a main effect of *Group* (F_6,997_=25.20, p≤0.001; μ^2^=0.13), with the observed disruptions in frontoparietal network connectivity increasing with diagnostic severity (Figure 3B; Table S3). Control B within-network connectivity was statistically similar (and even nominally higher) for the non-treatment seeking individuals with unipolar depression (0.61±0.16), as compared with healthy participants (0.58±0.16; p=0.82). Relative to the healthy participants, we observed reduced within-network control B connectivity in treatment-seeking patients with unipolar depression (0.51±0.14; p≤0.005), bipolar disorder without psychosis (0.44±0.16; p≤0.001), bipolar disorder with psychosis (0.42±0.18; p≤0.001), and the patients with schizophrenia (Group 1:0.46±0.18; Group 2: 0.35±0.16; ps≤0.001; See Table S3 for network interactions across each group). These analyses indicate that reports of disrupted frontoparietal network function in schizophrenia^23^ are replicable across recruitment sites. Further, they suggest that prior observations of frontoparietal network disruptions in psychosis reflect the presence of graded, transdiagnostic impairments in network connectivity evident in a host of Axis I pathologies.

### Evidence for both general and specific alterations in network connectivity

The observed reductions in functional connectivity were not specific to the frontoparietal network (Figure 2B; Table S2). When considering the remaining functional networks, with more than two parcels, for each network between 26 and 67 percent of associated within-network interactions exhibited a main effect of *Group* at the false discovery rate (q≤0.05) threshold. In order to explore the possibility of both diagnosis general and specific alterations in network function, mean network connectivity was computed for each patient group across every cortical network (Figure 4; Table S3). In treatment-seeking patients presenting with unipolar or bipolar depression without psychosis, impaired connectivity was localized to frontoparietal and limbic networks. Conversely, the presence of bipolar depression with psychosis and schizophrenia associated with a more generalized profile of reduced within-network connectivity throughout the cortex, encompassing the default network, the dorsal and ventral attention networks, and motor and visual systems. Below, we focus on the default network connectivity to highlight aspects of brain function that are preferentially disrupted in patients with psychotic illness, but not patients without psychotic symptoms. However, the focus on the default network should not be taken to imply that meaningful group or symptom specific properties are absent in other large-scale networks (see Figure 4).

**Figure 4.**
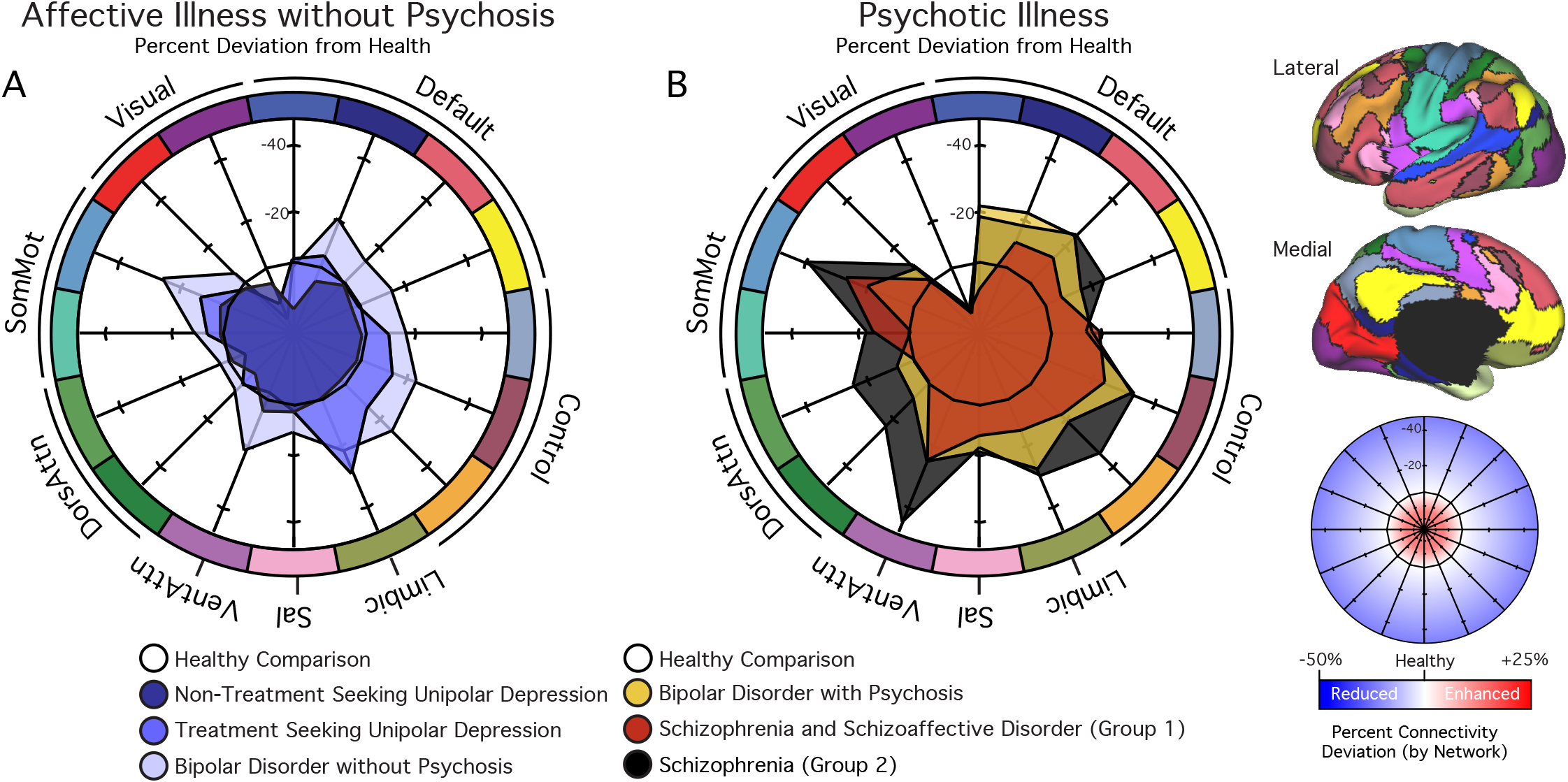
Polar plots display the percent difference in mean network connectivity for (A) patients with affective illnesses without psychosis or (B) psychotic illnesses relative to healthy comparison participants. The black hexadecagon in the center of each plot reflects the mean network correlations for the healthy comparison sample. Values outside the hexadecagon reflect decreased correlation strength for patients, relative to the healthy comparison sample. The graph scale reflects percent change values from healthy network connectivity (from −25 to 50 percent). See Figure 1 legend for explanation of abbreviations.

### Reduced default network connectivity in psychotic illnesses

The onset and maintenance of psychotic illness has been attributed to a breakdown in information processing, reflected in altered functional integration or connectivity across large-scale distributed brain networks. In line with prior work indicating broad disruptions across cortical association networks in psychosis, there is consistent evidence for aberrant default network connectivity in schizophrenia^16,23,36^. However, default network abnormalities have been observed across a range of psychiatric conditions, and it is not yet clear if this dysfunction represents a common feature of illness. We next examined whether default network disruption is linked to the presence of Axis 1 pathology, with and/or without psychotic features (Figure 5). Across default sub-networks A, B, and C we observed a main effect of diagnostic *Group* (Fs_2,1001_≥6.41, ps≤0.005; μ^2^s≥0.01; default D: F_2,1001_=2.66, p=0.07; μ^2^≤0.01). Follow-up tests revealed a global reduction in connectivity in patients with psychosis relative to both the non-psychotic illness (ps≤0.01) and healthy comparison samples (ps≤0.001; Table S2). Default network connectivity did not significantly differ between the healthy comparison and the affective illness without psychosis group (ps≥0.06)

**Figure 5.**
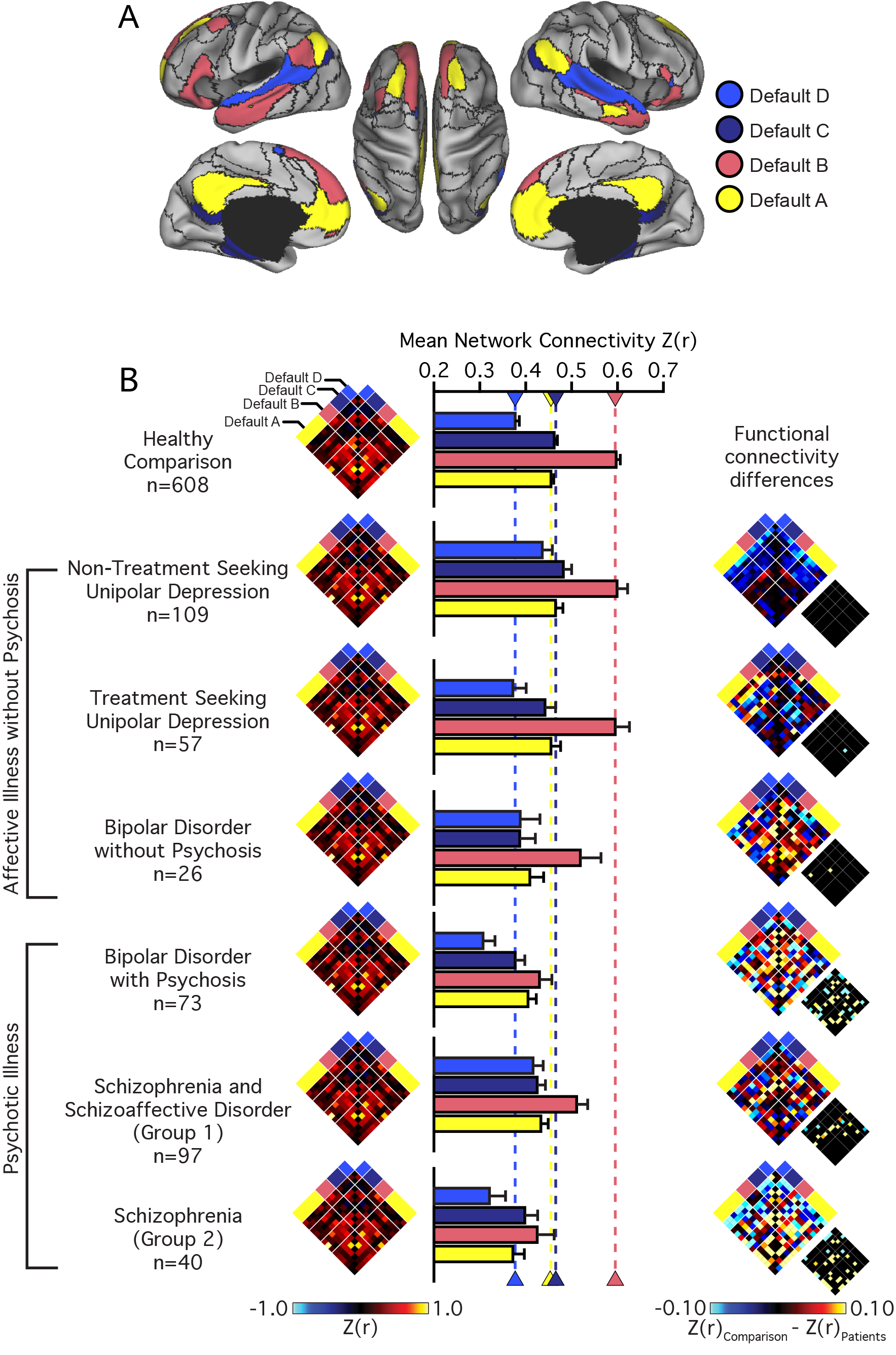
Psychotic illness associates with a preferential reduction in default network integrity. (A) The colored aspects of cortex reflect regions estimated to be within the A, B, C, and D aspects of the default network. Black lines denote network boundaries defined through the 17-network solution from Yeo et al. 2011. (B) Functional connectivity matrices for the 14 left and right hemisphere regions of the default network shown for the healthy comparison and patient groups. Bar graphs reflect mean network connectivity for each group within default A, B, C, and D. Error bars denote standard error. Functional connectivity difference matrices were obtained by an analysis of variance of z-transformed Pearson correlation values after linear regression of the effects of coil, scanner, console software version, age, sex, race, ethnicity, and handedness. Differences significant at false discovery rate (q≤0.05) are shown in each panel just to the lower right of the unthresholded matrix.

To further explore the nature of functional connectivity changes in the default network, we conducted additional analyses across individual DSM-IV diagnoses. Analyses of region-to-region correlation strength across the default network revealed a main effect of *Group* across each sub-network (A-D; Fs_6,997_≥3.45, ps≤0.005; μ^2^s≥0.02; Table S3). We observed an increase in default D connectivity for the non-treatment seeking individuals with unipolar depression (0.44±0.27) relative to the healthy comparison sample (0.38±0.22; p≤0.005). With the exception of a subtle decrease in default C connectivity for the patients with bipolar disorder (p≤0.05), no other effects of diagnoses were observed for the patients without psychosis relative to the healthy comparison sample (ps≥0.09). Global decreases in default network connectivity were observed in the patients diagnosed with bipolar disorder with psychosis (ps≤0.01). Relative to the healthy comparison participants, reduced default B and C within-network connectivity was observed in the patients with schizophrenia (ps≤0.05; See Table S3 for network interactions across each group). Collectively, these results indicate that while altered patterns of frontoparietal connectivity may be shared across diagnostic groups, perturbations within other large-scale networks may reflect the expression of specific symptoms or presence or certain diagnostic groups.

## DISCUSSION

The present analyses reveal the existence of functional connectomic profiles that bridge diagnostic categories, aligning with clinical symptoms in a graded fashion. Here, we identified a pattern of disrupted connectivity within the frontoparietal network that was evident across specific categories of impairment (psychosis) and clinical diagnoses (unipolar depression, bipolar disorder, and schizophrenia or schizoaffective disorder). Suggesting that these alterations in connectivity may track the severity of illness, frontoparietal network impairments were observed in treatment-seeking patients with unipolar depression recruited from inpatient and partial hospital programs, but not those recruited from the general community who were not treatment-seeking. Highlighting the presence of diagnostic and symptom specific profiles of connectivity, aspects of the default network exhibited reduced connectivity in patients with psychotic illness, a pattern of impaired network functioning that was absent in patients with unipolar depression or bipolar disorder without psychosis. Collectively, these observations suggest that complex psychiatric symptoms are associated with specific patterns of abnormal connectivity across the connectome, with the disturbance of individual systems preferentially contributing to certain symptom domains, which can present in disorder-general manner.

Our findings of transdiagnostic disruptions in frontoparietal network connectivity are consistent with prior work in both schizophrenia^37–39^, unipolar depression^27,40,41^, and bipolar disorder^42,43^, where there is converging evidence for abnormalities in cognitive control and context processing^8,21^. By studying multiple patient populations simultaneously, without prejudice toward ascertainment or diagnostic label, our findings allow for the simultaneous characterization and comparison of psychiatric connectomes across both affective and psychotic illnesses. Further, while much of the prior work in this domain has focused on circumscribed profiles of dysfunction in either dorsolateral prefrontal or anterior cingulate cortices, here we provide evidence indicating broad frontoparietal network impairments that span aspects of frontal, parietal, temporal, and medial prefrontal components of this network. Although these data support the view that the frontoparietal network may underlie a diverse set of cognitive processes impaired in multiple disorders, one outstanding question centers on the extent to which impaired frontoparietal connectivity in individual patients may reflect a primary factor associated with disease onset and maintenance, or a secondary consequence of illness^8^. Suggesting the likely presence of pre-illness shifts in network connectivity, prefrontal dysfunctions related to context processing have been identified in never-medicated patients with schizophrenia early in the course of the illness^37^. Highlighting a degree of symptom specificity, in a study of psychotic patients with and without cognitive dysfunction, we recently discovered that distinct frontoparietal subnetworks may link to cognitive capacity (i.e., control A subnetwork) and psychiatric symptoms (i.e., control B subnetwork), respectively^44^.

A considerable body of evidence has accumulated over recent years suggesting that the presence of altered default network functioning may mark psychotic illness^45^. Encompassing aspects of ventral and dorsal medial prefrontal, posterior/retrosplenial, and inferior parietal cortices, the default network is hypothesized to underpin self-referential processing and principal aspects of mental simulation^46^. Core symptoms of psychotic illness arise from misattributions of thought and a blurring of the boundaries separating internal cognition from the external world^46,47^. Consistent with our reported analyses, these converging lines of evidence suggest an association linking impaired default network functioning and the occurrence of psychotic symptoms (e.g., hallucinations, delusions, and thought distortions). Critically however, the present results should not be taken to suggest that default network disruption is specific to patients with psychotic illness. Indeed, in line with prior reports^48^, a muted decrease in default network functioning was observed in patients with bipolar disorder without psychosis. Rather, our data support the view that default network functions may underlie a set of cognitive processes impaired in multiple disorders^45^, with variability in network-level functioning linking with corresponding network-associated symptom expression^21^. Patients with unipolar depression, for example, have been found to display aberrant intrinsic connectivity^49^ as well as heightened stimulus-induced activity in aspects of the default network while viewing and reappraising negative images^50^. These data are consistent with our observation of increased default D subnetwork connectivity in non-treatment seeking individuals with depression, a functional profile that was absent in treatment-seeking patients recruited from clinical units. Given this observed variability within diagnoses, future high-throughput data collection efforts will be necessary to establish the manner in which individual-specific connectome architecture might serve as a dimensional fingerprint of human behavior, predicting symptom profiles in patient populations with varying degrees of clinical severity.

Analyses that link functional connectomes to individual differences in behavior, symptom profiles, and severity of illness represent a tremendous opportunity for the field. While it may not be feasible to identify isolated features of brain biology that cleanly distinguish populations of patients with psychiatric illness, multivariate fingerprints of pathology may eventually emerge. To establish such points of separation, our data collection and analytic efforts need to incorporate dimensional measures of clinical severity across a broad range of patient populations, recruited from diverse clinical settings at varying phases of illness. In order to generate such high-dimensional datasets we will need to reassess our current scientific approach, extending beyond conventional clinic- or research laboratory specific collection efforts. Our current analyses reflect the combined efforts of multiple research groups collaborating to collect data with a harmonized acquisition sequence^30^. This cross-lab collaborative effort allowed us to partially disentangle the relations between clinical diagnoses and degree of treatment seeking.

Readers should note that there are limitations on the conclusions that can be drawn from the current analyses. Given the cross-lab collaborative nature of the present work, a consistent self-report battery was not available for analysis across the included participants. Accordingly, we are unable to make claims regarding associations that may link network function with the presence, absence, or severity of specific dimensional symptom profiles. To make progress in this domain, our clinical recruitment strategies and analytic efforts will need to coordinate across research labs to standardize imaging acquisition protocols as well as dense demographic, symptom, and behavioral batteries^1^. Additionally, while we can compare and contrast treatment-seeking and non-treatment seeking environments individuals with unipolar depression, we are unable to account for other factors that may contribute to access to care and utilization. For instance, the non-treatment seeking individuals with depression were not currently taking psychiatric medications. This is in contrast to individuals already in treatment who were prescribed varying forms of psychiatric medication. Additionally, internalized and treatment stigma can associate with reduced help-seeking in some patient populations^51^. Consequently, despite evaluating over 1,000 individuals, we are limited in the conclusions we can draw when comparing and contrasting the connectomic profiles observed in non-treatment seeking individuals recruited from the community and populations of patients recruited from partial and inpatient hospitalization programs. Smaller scale longitudinal studies suggest the presence of abnormal functional connectivity in individuals presenting to the emergency room seeking care for schizophrenia or related diagnoses, even prior to starting psychiatric medication^52^. Moreover, connectomic changes are apparent in early phases of antipsychotic treatment^53^. This literature suggests that differences in symptom severity – rather than medication per se – may underlie the extent and degree of changes observed in our present analyses. However, future longitudinal research designs will be critical for fully disentangling the effects of treatment, fluctuating symptom-severity, and illness course on brain function.

The unprecedented growth of big data in neuroscience provides opportunities for researchers seeking to understand how brain functions influence suites of behaviors and associated illness risk. In the present analyses we make use of a large resource of individuals with imaging data, spanning domains of psychopathology, levels of acuity, and engagement with care. This heterogeneous sample of participants represented a broad range of symptom profiles and illness severity, including individuals with selfreported mental health, non-treatment seeking forms of depression, and treatment-seeking forms of unipolar depression, bipolar disorder, and severe psychotic illness. Our analyses revealed aspects of the frontoparietal control network that are commonly disrupted across diagnostically distinct forms of severe pathology, whether psychotic or non-psychotic affective in nature. Here, as well, we established both shared and unique functional alterations in affective and psychotic illnesses. For instance, a preferential reduction in default network integrity was evident in patients with psychotic illness, but absent in affective illnesses without psychosis. These analyses highlight the potential to discover individualized network profiles that are predictive of symptom-relevant cognitive domains, both within and across diagnostic boundaries, as exemplified in the B-SNIP effort^54^ and our own ongoing work. In conclusion, this study provides a novel and comprehensive characterization of connectomic dysfunction in a range of psychopathological conditions that matches well with the core deficits observed in these populations. These data have important implications for the future creation of connectome-based models that predict behavior, an approach with the potential to account for symptom comorbidity while simultaneously explaining the biological process that give rise to the diversity of clinical presentations.

## METHODS

Between November 2008 and June 2017, functional magnetic resonance imaging (fMRI) data were collected from a total of 1,010 individuals, including 210 diagnosed with a primary psychotic disorder (137 meeting criteria for schizophrenia or schizoaffective disorder, 73 with bipolar disorder with psychosis), 192 presenting with a primary affective disorder without psychosis (26 with bipolar disorder without psychosis, 57 treatment seeking individuals with unipolar depression, 109 non-treatment seeking individuals with unipolar depression), and 608 demographically and data-quality matched healthy comparison participants recruited through an ongoing, large-scale study of brain imaging and genetics^30^. Details regarding participant recruitment and characterization, as well as the demographic and clinical characteristics of the patient and matched healthy comparison samples are available in Supplemental Table S1. In brief, patients were recruited from clinical services at Massachusetts General Hospital (MGH) or McLean Hospital through the procedures detailed in Baker et al., 2014^23^. Nontreatment seeking individuals who met diagnostic criteria for unipolar depression were recruited from the surrounding Boston area using the procedures detailed in Dillon et al., 2014^55^. Participants provided written informed consent in accordance with guidelines set by the Partners HealthCare Institutional Review Board and the Harvard University Committee on the Use of Human Subjects in Research.

### MRI data acquisition

Imaging data were collected on 3T Tim Trio scanners (Siemens) using either 12- or 32-channel phased-array head coils at Harvard University, MGH, or McLean Hospital as detailed in Holmes et al., 2015^30^. Briefly, structural data included a high-resolution, multi-echo T1-weighted magnetization-prepared gradient-echo image (144 slices, TR=2200ms, TI=1100ms, TE=1.54ms for image 1-7.01ms for image 4, flip angle=7°, voxels=1.2mm^3^, FOV=230). Functional data were acquired using a gradient-echo echoplanar imaging sequence (47 axial slices, interleaved with no gap), 124 time points, repetition time=3000ms, echo time=30ms, flip angle=85°, voxels=3mm^3^, FOV=216). Participants were instructed to remain still and keep their eyes open, while blinking normally. Although no fixation image was used, participants with psychotic illness were monitored via eye tracking video to ensure compliance during functional scans. Software upgrades (VB13, VB15, VB17) occurred during data collection. All results are reported after partialing out variance associated with coil, scanner (Harvard Bay 1, McLean Bay 1, MGH Bay 4, MGH Bay 8, etc.), and software upgrade, as well as age, sex, handedness, race, and ethnicity.

### Preprocessing

Data were analyzed with a series of steps common to intrinsic connectivity analyses^31–33^ and further elaborated in Holmes et al., 2015^30^ and Yeo et al., 2011^34^. Preprocessing included discarding the first four volumes of each run to allow for T1-equilibration effects, compensating for slice acquisition-dependent time shifts per volume, and correcting for head motion using rigid body translation and rotation. Additional steps involved the removal of constant offset and linear trends over each run and the use of a temporal filter to retain frequencies below 0.08 Hz. Sources of spurious variance, along with their temporal derivatives, were removed through linear regression. These included six parameters obtained by correction for rigid body head motion, the signal averaged over the whole brain, the signal averaged over the ventricles, and the signal averaged over the deep cerebral white matter. Functional data were first aligned to the structural image using the FreeSurfer software package, smoothed using a 6mm kernel applied in surface space, and down-sampled to a 4mm mesh (Yeo et al. 2011).

### Functional parcellation

Cortical functional coupling matrices were computed for each participant, across all available regions within the 17 network functional parcellation of Yeo et al. 2011^34^ (Figure 1A). This parcellation consisted of 122 cortical regions composed of 61 roughly symmetric territories in the left and right hemispheres^23^. Correlation matrices were constructed to include all regional pairs arranged by network membership. Pearson correlation coefficients were computed between each regional fMRI time course, averaged across all vertices within the region, and the mean fMRI time course for every other region (Figure 1B-D). Correlation values were z-transformed to increase normality of the correlation distribution and compared across groups using an analysis of variance after linear regression of nuisance variables. Reported tests survived correction for multiple comparisons using a family-wise error rate (FWER; Bonferroni procedure) of p≤0.05 or false discovery rate (FDR) of q≤0.05.

## Supporting information

## ACKNOWLEDGEMENTS

This work was supported by the Brain & Behavior Research Foundation (A.J.H and D.G.D.), the Taplin Family Foundation (D.Ö.), and the National Institute of Mental Health (Grant K23MH104515 to J.T.B., Grants F32MH081394 and R00MH094438 to D.G.D., Grants R37MH068376 and 1R01MH101521 to D.A.P., Grant K23MH100623 to R.O.B., Grants K23MH079982-01A1 and R01MH094594 to D.Ö, and Grant K01MH099232 to A.J.H.). Analyses were made possible by the resources provided through Shared Instrumentation Grants 1S10RR023043 and 1S10RR023401. Data were provided in part by the Brain Genomics Superstruct Project of Harvard University and Massachusetts General Hospital (Principal Investigators: Randy L. Buckner, Jordan W. Smoller, and J.L.R.) with support from the Center for Brain Science Neuroinformatics Research Group, the Athinoula A. Martinos Center for Biomedical Imaging, and the Center for Genomic Medicine. Both the Simons Foundation and the Howard Hughes Medical Institute supported Randy L. Buckner’s work on the GSP, and we are immensely grateful for their invaluable contributions on that project.

