## Supplementary material for "Functional connectomics of affective and psychotic pathology"

Table 1. Participant demographics

| | Healthy Comparison | Non-Treatment Seeking Depression | Treatment Seeking Depression | Bipolar Disorder (w/o psychosis) | Bipolar Disorder (psychosis) | Scz Group1 | Scz Group 2 | | | Healthy Comparison vs | | | | | |
|---|---|---|---|---|---|---|---|---|---|---|---|---|---|---|---|
| | | | | | | | | | | NTSD | TSD | BD w/o | BD w/ | SZ gp1 | SZ gp2 |
| Network | n=608 | n=109 | n=57 | n=26 | n=73 | n=97 | n=40 | $F_{6,1003}/\chi^2$ | p-value | p-values | | | | | |
| Age | 33.6±13.5 | 30.1±10.9 | 35.0±13.7 | 38.7±15.3 | 31.3±11.4 | 33.3±12.0 | 44.8±10.6 | 1248.6 | <0.001 | | | | | | **** |
| Age range | 18-71 | 18-64 | 18-68 | 20-71 | 18-63 | 18-65 | 20-62 | | | | | | | | |
| Percent female | 44.9 | 66.1 | 57.9 | 53.8 | 35.6 | 32.0 | 22.5 | 41.5 | <0.001 | **** | | | | * | *** |
| Percent right handed | 91.4 | 100.0 | 89.5 | 92.3 | 84.9 | 84.5 | 80.0 | 30.3 | <0.005 | ** | | | | * | *** |
| Percent White (not Hispanic or Latino) | 65.1 | 65.2 | 80.4 | 69.2 | 75.3 | 70.1 | 67.5 | 4.5 | 0.62 | | | | | | |
| Slice SNR | 175.3±51.2 | 196.1±67.3 | 195.6±53.3 | 177.6±40.4 | 140.7±65.4 | 166.2±66.2 | 144.3±67.0 | 10.7 | <0.001 | ** | | | **** | | * |
| Number of micro-movements | 20.3±24.7 | 15.1±20.4 | 24.5±23.1 | 24.8±17.4 | 35.1±31.8 | 26.9±26.6 | 37.9±35.6 | 8.4 | <0.001 | | | | **** | | **** |

*Note: NTSD: community non-treatment seeking depression; TSD: unipolar treatment seeking depression; BD w/o: bipolar disorder without psychosis; BD w/: bipolar disorder with psychosis; SZ gp1: schizophrenia group 1; SZ gp2: schizophrenia group 2. Micro-movements reflect the number of relative translations in 3D space ≥0.1 mm. *P*-values for group comparisons * ≤0.05 **≤0.01 *** ≤0.005 **** ≤0.001

Supplemental Table 1 Cont. Participant demographics

| | Non-Treatment Seeking Depression vs | | | | | Treatment Seeking Depression vs | | | | Bipolar Disorder (w/o psychosis) vs | | | Bipolar Disorder (psychosis) vs | | Scz Group1 vs Scz Group2 |
|---|---|---|---|---|---|---|---|---|---|---|---|---|---|---|---|
| | TSD | BD w/o | BD w/ | SZ gp1 | SZ gp2 | BD w/o | BD w/ | SZ gp1 | SZ gp2 | BD w/ | SZ gp1 | SZ gp2 | SZ gp1 | SZ gp2 | |
| Network | p-values | | | | | p-values | | | | p-values | | | p-values | | p-values |
| Age | | | * | | **** | | | | *** | | | | | **** | **** |
| Age range | | | | | | | | | | | | | | | |
| Percent female | | | **** | **** | **** | | * | *** | **** | | * | ** | | | |
| Percent right handed | *** | *** | **** | **** | **** | | | | | | | | | | |
| Percent White (not Hispanic or Latino) | | | | | | | | | | | | | | | |
| Slice SNR | | | **** | *** | **** | | **** | * | **** | * | | | * | | |
| Number of micro-movements | | | **** | * | **** | | | | | | | | | | |

*Note: NTSD: community non-treatment seeking depression; TSD: unipolar treatment seeking depression; BD w/o: bipolar disorder without psychosis; BD w/: bipolar disorder with psychosis; SZ gp1: schizophrenia group 1; SZ gp2: schizophrenia group 2. *P*-values for group comparisons * ≤0.05 **≤0.01 *** ≤0.005 **** ≤0.001

Supplemental Table 2. Network interactions affected by psychiatric illnesses with or without psychosis

| | Healthy Comparison | Patients without psychosis | Patients with psychosis | | | | HC vs Pw/o | HC vs PP | PP vs Pw/o |
|---|---|---|---|---|---|---|---|---|---|
| Network | n=608 | n=192 | n=210 | $F_{2,1001}$ | $\mu^2$ | p-value | p-values | | |
| Default D | 0.38±0.22 | 0.41±0.24 | 0.35±0.21 | 2.66 | 0.01 | 0.07 | 0.06 | 0.33 | ≤0.05 |
| Default C | 0.46±0.19 | 0.46±0.17 | 0.40±0.16 | 8.84 | 0.02 | ≤0.001 | 0.84 | ≤0.001 | ≤0.005 |
| Default B | 0.60±0.23 | 0.60±0.25 | 0.45±0.28 | 23.72 | 0.05 | ≤0.001 | 0.55 | ≤0.001 | ≤0.001 |
| Default A | 0.46±0.15 | 0.46±0.16 | 0.40±0.18 | 6.41 | 0.01 | ≤0.005 | 0.93 | ≤0.001 | ≤0.01 |
| Control C | 0.66±0.22 | 0.64±0.24 | 0.55±0.19 | 14.42 | 0.03 | ≤0.001 | 0.14 | ≤0.001 | ≤0.005 |
| Control B | 0.58±0.16 | 0.56±0.16 | 0.42±0.18 | 63.95 | 0.11 | ≤0.001 | ≤0.01 | ≤0.001 | ≤0.001 |
| Control A | 0.33±0.11 | 0.31±0.11 | 0.26±0.11 | 25.65 | 0.05 | ≤0.001 | 0.06 | ≤0.001 | ≤0.001 |
| Limbic | 0.38±0.26 | 0.33±0.28 | 0.31±0.27 | 6.52 | 0.01 | ≤0.005 | 0.08 | ≤0.001 | 0.21 |
| Salience | 0.45±0.13 | 0.45±0.13 | 0.38±0.14 | 20.39 | 0.04 | ≤0.001 | 0.18 | ≤0.001 | ≤0.001 |
| Ventral attention | 0.46±0.14 | 0.46±0.15 | 0.40±0.16 | 9.86 | 0.02 | ≤0.001 | 0.48 | ≤0.001 | ≤0.005 |
| Dorsal attention B | 0.48±0.17 | 0.50±0.17 | 0.45±0.19 | 3.01 | 0.01 | ≤0.05 | 0.35 | ≤0.05 | ≤0.05 |
| Dorsal attention A | 0.68±0.23 | 0.71±0.22 | 0.63±0.27 | 4.54 | 0.01 | ≤0.05 | 0.24 | ≤0.05 | ≤0.005 |
| Somatomotor B | 0.67±0.17 | 0.65±0.18 | 0.62±0.17 | 6.88 | 0.01 | ≤0.001 | 0.09 | ≤0.001 | 0.16 |
| Somatomotor A | 0.61±0.25 | 0.58±0.26 | 0.44±0.26 | 26.58 | 0.05 | ≤0.001 | ≤0.05 | ≤0.001 | ≤0.001 |
| Visual peripheral | 1.05±0.25 | 1.08±0.26 | 0.99±0.24 | 2.68 | 0.01 | 0.07 | 0.53 | ≤0.05 | ≤0.05 |
| Visual central | 0.72±0.31 | 0.80±0.32 | 0.84±0.27 | 14.83 | 0.03 | ≤0.001 | ≤0.01 | ≤0.001 | 0.08 |

*Note: Values reflect mean of raw z-transformed Pearson correlation values ± standard deviation across all regional interactions for that network; HC: healthy comparison participants; Pw/o: patients without psychosis; PP: patients with psychosis.

Supplemental Table 3. Network interactions affected by psychiatric illnesses with or without psychosis

| | Healthy Comparison | Non-Treatment Seeking Depression | Treatment Seeking Depression | Bipolar Disorder (w/o psychosis) | Bipolar Disorder (psychosis) | Scz Group1 | Scz Group 2 | | | | Healthy Comparison vs | | | | | |
|---|---|---|---|---|---|---|---|---|---|---|---|---|---|---|---|---|
| | | | | | | | | | | | NTSD | TSD | BD w/o | BD w/ | SZ gp1 | SZ gp2 |
| Network | n=608 | n=109 | n=57 | n=26 | n=73 | n=97 | n=40 | $F_{6,997}$ | $\mu^2$ | p-value | p-values | | | | | |
| Default D | 0.38±0.22 | 0.44±0.27 | 0.37±0.20 | 0.37±0.20 | 0.31±0.20 | 0.41±0.22 | 0.29±0.17 | 3.67 | 0.02 | ≤0.001 | *** | | | ** | | |
| Default C | 0.46±0.19 | 0.48±0.18 | 0.45±0.15 | 0.38±0.14 | 0.38±0.14 | 0.42±0.17 | 0.36±0.14 | 4.98 | 0.03 | ≤0.001 | | | * | **** | * | * |
| Default B | 0.60±0.23 | 0.63±0.25 | 0.59±0.25 | 0.49±0.22 | 0.44±0.26 | 0.51±0.28 | 0.35±0.30 | 9.47 | 0.05 | ≤0.001 | | | | **** | **** | **** |
| Default A | 0.46±0.15 | 0.48±0.16 | 0.46±0.14 | 0.40±0.16 | 0.41±0.17 | 0.43±0.19 | 0.34±0.16 | 3.45 | 0.02 | ≤0.005 | | | | ** | | **** |
| Control C | 0.66±0.22 | 0.68±0.25 | 0.60±0.22 | 0.55±0.21 | 0.57±0.20 | 0.55±0.19 | 0.54±0.17 | 5.79 | 0.03 | ≤0.001 | | | | **** | **** | * |
| Control B | 0.58±0.16 | 0.61±0.16 | 0.51±0.14 | 0.44±0.16 | 0.42±0.18 | 0.46±0.18 | 0.35±0.16 | 25.20 | 0.13 | ≤0.001 | | *** | **** | **** | **** | **** |
| Control A | 0.33±0.11 | 0.34±0.11 | 0.29±0.10 | 0.25±0.08 | 0.28±0.11 | 0.28±0.11 | 0.20±0.08 | 12.71 | 0.07 | ≤0.001 | | * | **** | **** | **** | **** |
| Limbic | 0.38±0.26 | 0.36±0.28 | 0.27±0.28 | 0.29±0.24 | 0.30±0.25 | 0.35±0.27 | 0.26±0.28 | 3.63 | 0.02 | ≤0.001 | | ** | | *** | | * |
| Salience | 0.45±0.13 | 0.46±0.12 | 0.44±0.12 | 0.39±0.14 | 0.38±0.14 | 0.40±0.14 | 0.34±0.15 | 7.52 | 0.04 | ≤0.001 | | | | **** | **** | **** |
| Ventral attention | 0.46±0.14 | 0.48±0.16 | 0.46±0.14 | 0.41±0.14 | 0.42±0.15 | 0.41±0.17 | 0.33±0.14 | 4.09 | 0.02 | ≤0.001 | | | | * | ** | **** |
| Dorsal attention B | 0.48±0.17 | 0.52±0.18 | 0.48±0.15 | 0.45±0.18 | 0.45±0.16 | 0.48±0.21 | 0.35±0.15 | 2.81 | 0.02 | ≤0.01 | | | | | | **** |
| Dorsal attention A | 0.68±0.23 | 0.72±0.23 | 0.73±0.20 | 0.64±0.17 | 0.64±0.26 | 0.68±0.26 | 0.46±0.23 | 4.15 | 0.02 | ≤0.001 | | | | | | **** |
| Somatomotor B | 0.67±0.17 | 0.68±0.20 | 0.64±0.15 | 0.59±0.17 | 0.68±0.18 | 0.59±0.16 | 0.55±0.14 | 4.95 | 0.03 | ≤0.001 | | | | | **** | **** |
| Somatomotor A | 0.61±0.25 | 0.64±0.27 | 0.53±0.21 | 0.44±0.25 | 0.50±0.25 | 0.45±0.25 | 0.32±0.25 | 10.94 | 0.06 | ≤0.001 | | | *** | **** | **** | **** |
| Visual peripheral | 1.05±0.25 | 1.10±0.30 | 1.07±0.20 | 0.99±0.25 | 0.99±0.24 | 1.03±0.24 | 0.89±0.21 | 1.77 | 0.01 | 0.10 | | | | * | | * |
| Visual central | 0.72±0.31 | 0.76±0.34 | 0.83±0.29 | 0.87±0.26 | 0.85±0.21 | 0.84±0.32 | 0.81±0.21 | 5.65 | 0.03 | ≤0.001 | | * | * | **** | **** | * |

*Note: Values reflect mean of raw z-transformed Pearson correlation values ± standard deviation across all regional interactions for that network. NTSD: community non-treatment seeking depression; TSD: unipolar treatment seeking depression; BD w/o: bipolar disorder without psychosis; BD w/: bipolar disorder with psychosis; SZ gp1: schizophrenia group 1; SZ gp2: schizophrenia group 2. *P*-values for group comparisons * ≤0.05 **≤0.01 *** ≤0.005 **** ≤0.001

Supplemental Table 3 Cont. Network interactions affected by psychiatric illnesses with or without psychosis

| | Non-Treatment Seeking Depression vs | | | | | Treatment Seeking Depression vs | | | | Bipolar Disorder (w/o psychosis) vs | | | Bipolar Disorder (psychosis) vs | | Scz Group1 vs Scz Group2 |
|---|---|---|---|---|---|---|---|---|---|---|---|---|---|---|---|
| | TSD | Bw/o | B | S1 | S2 | Bw/o | B | S1 | S2 | B | S1 | S2 | S1 | S2 | |
| Network | p-values | | | | | p-values | | | | p-values | | | p-values | | p-values |
| Default D | * | | **** | | *** | | | | | | | | **** | | * |
| Default C | | *** | **** | ** | *** | | * | | | | | | | | |
| Default B | | | **** | * | **** | | *** | * | **** | | | | * | | * |
| Default A | | | ** | | **** | | | | * | | | | | | * |
| Control C | | * | *** | **** | * | | | | | | | | | | |
| Control B | * | **** | **** | **** | **** | | **** | * | **** | | | | * | | * |
| Control A | ** | **** | **** | **** | **** | | | | **** | | | | | * | *** |
| Limbic | * | | * | | * | | | | | | | | | | |
| Salience | | | **** | * | * | | **** | * | * | | | | | | |
| Ventral attention | | | | | ** | | * | * | * | | | | | | |
| Dorsal attention B | | | * | | **** | | | | ** | | | | | | *** |
| Dorsal attention A | | | | | **** | | | | **** | | | * | | * | **** |
| Somatomotor B | | | | * | * | | | | | | | | **** | **** | |
| Somatomotor A | | * | | **** | **** | | | * | **** | | | | | *** | * |
| Visual peripheral | | | * | | * | | * | | * | | | | | | |
| Visual central | | | * | * | | | | | | | | | | | |

*Note: CD: community depression; UD: unipolar depression; Bw/o: bipolar depression without psychosis; B: bipolar depression with psychosis; S1: schizophrenia group 1; S2: schizophrenia group 2. P-values for group comparisons * ≤0.05 **≤0.01 *** ≤0.005 **** ≤0.001
